# A probabilistic framework to dissect functional cell-type-specific regulatory elements and risk loci underlying the genetics of complex traits

**DOI:** 10.1101/059345

**Authors:** Yue Li, Jose Davila-Velderrain, Manolis Kellis

## Abstract

Dissecting the physiological circuitry underlying diverse human complex traits associated with heritable common mutations is an ongoing effort. The primary challenge involves identifying the relevant cell types and the causal variants among the vast majority of the associated mutations in the noncoding regions. To address this challenge, we developed an efficient probabilistic framework. First, we propose a sparse group-guided learning algorithm to infer cell-type-specific enrichments. Second, we propose a fine-mapping Bayesian model that incorporates as Bayesian priors the sparse enrichments to infer risk variants. Using the proposed framework to analyze 32 complex human traits revealed meaningful tissue-specific epigenomic enrichments indicative of the relevant disease pathologies. The prioritized variants exhibit prominent tissue-specific epigenomic signatures and significant enrichments for eQTL and conserved elements. Together, we demonstrate the general benefits of the proposed integrative framework in elucidating meaningful tissue-specific epigenomic elements from large-scale correlated annotations and the implicated functional variants for future experimental interrogation.

## 1 Introduction

Over the past several years, genome wide association studies (GWAS) have identified genetic signals for many complex human traits, that exhibit intricate polygenic architectures and comorbidities among similar phenotypes. Systematic investigation of these GWAS data involves not only identifying the associated variants but also the relevant cell-types, and it promises to decipher the underlying epidemiology at the single-nucleotide resolution ^1–4^.

However, there are several challenges in this approach that hinder further progress. These include (1) difficulty in interpreting the function of variants in the noncoding regions, which contribute about 90% of the current GWAS catalog ^5^; (2) the lack of statistical power to detect causal SNPs that are under-represented in the population due to low minor allele frequency and thus exhibit small effect size ^6^; (3) insufficient knowledge about the potentially relevant cell types and tissues that are disrupted by the mutations; (4) the fact that the true causal variants are often harbored within large haplotype blocks spanning many kilobases of the human genome, where SNPs are correlated with one another via linkage disequilibrium (LD)^1^.

To help interpret non-coding elements and gain insights into their potential regulatory functions, several international consortia have recently released a plethora of genome-wide reference annotations. In particular, the ENCODE/Roadmap Epigenomics consortium generated and made publicly available epigenomic annotations in diverse cell types and tissues, providing a multi-dimensional reference map to elucidate enhancer/repressor locations indicative of cis-regulatory functions^8^. The 1000 Genome consortium provided genotype information of hundreds of individuals that share ancestral haplotypes, which can be used to estimate LD structure in terms of SNP-by-SNP genetic correlation (as a surrogate to the in-sample LD from the GWAS cohort that are often unavailable) to help disentangle functional signals from those stemming from co-inherited but passenger SNPs occurring within the same haplotypes ^9^.

Several studies have implicated enrichment of GWAS variants in putative regulatory elements - e.g., enhancer-associated histone modifications, regions of open chromatin, and conserved non-coding elements^3,10–13^ ^14,15^. Moreover, this overrepresentation has also been used to predict relevant cell types and non-coding annotations for specific traits^16^. Several recently developed methods are able to leverage the GWAS summary statistics in terms of marginal statistics of the association of each SNP with the trait of interest ^17,18^ as well as functional annotations to aid the inference of risk variants ^19–23^. However, it remains a challenge to explain genetic signals and infer causal variants based on *genome-wide* functional and tissue-specific enrichments. The hundreds of epigenomic annotations, although informative, are frequently correlated, making the interpretation of their connections with the causal variants difficult. Specifically, many annotations overlap with one another due to the sharing of regulatory elements in the genome and across tissues. Thus, it is challenging to interpret the tissue-specific enrichments when simultaneously using all of the annotations to explain the genetic signals and pinpoint the potential tissues of action.

In this article, we describe a novel probabilistic framework to infer sparse interpretable genome-wide functional enrichments as well as the locus-specific influence of these annotations to ultimately infer risk variants potentially underlying the genetic association through tissue specific perturbations. We apply our proposed method to investigate 32 complex human traits using the publicly available full summary statistics each containing 5-11 million genotyped or imputed SNPs. To leverage the evidence of tissue-specific epigenomic activities, we harness a large compendium of functional genomic and epigenomic annotations precompiled from ENCODE/Roadmap consortium, including four epigenomic marks implicated in promoting transcription (H3K4me1, H3K4me3, H3K27ac, H3K9ac) across 100 well-characterized cell types and tissues^8^. Because of the interpretability of the proposed model, we are able to make intuitive biological observations based on our inference results, allowing us to revisit these many complex human traits from a novel system-level perspective. Our method is implemented as an R package that can be easily apply to analyzing other complex traits.

## 2 Results

### 2.1 A novel probabilistic framework to infer interpretable genome-wide tissue-specific enrichment patterns and locus-specific risk variants

To infer relevant cell types and variants associated with each trait, we take into account three lines of evidence (**Fig. 1a**): (1) genome-wide genetic signals, in terms of marginal summary statistics of each SNP; (2) functional genomic and epigenomic reference annotations overlapping the SNPs; and (3) linkage disequilibrium, by estimating SNP-by-SNP Pearson correlation from the 1000 Genome reference panel. The general scheme of our learning algorithm follows an expectation-maximization (EM) formulation: at E-step, we infer the posterior probabilities of causal SNPs by fixing their functional enrichments; at M-step we learn the functional enrichments over all of the annotations in concordance to the posterior probabilities of the SNP associations. We then alternate between the E and M-step until some convergence criterion is satisfied. Here the SNP posteriors are defined by the *likelihood* dictated by the genetic signals and the *prior* dictated by a logistic function, which is a weighted linear combination of the annotations. Because of the linearity of the model, we interpret the highly positive coefficient weights as *functional enrichments* of the corresponding annotations.

**Figure 1.**
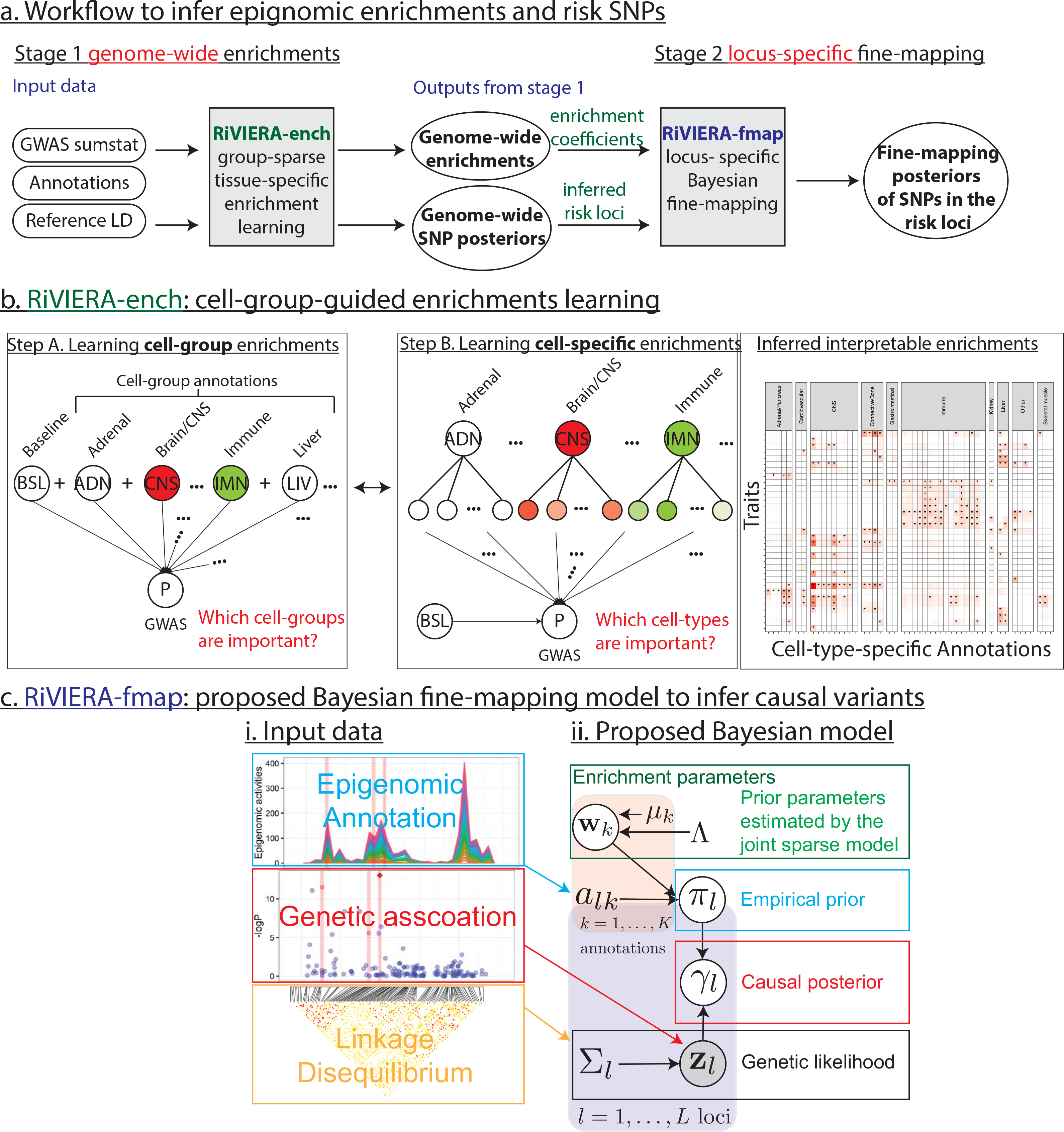
Integrative approach to infer tissue-specific enrichments and risk mutations. (a) Workflow to infer epigenomic enrichments and risk SNPs. Taking as inputs GWAS summary statistics, functional reference annotations, and reference LD, RiVIERA-glass infers the genome-wide tissue-specific enrichments in a sparse and interpretable fashion. The output enrichments are then incorporated into the Bayesian priors of the fine-mapping RiVIERA-fmap model. The final outputs are locus-specific the posterior probabilities of the SNPs. (b) Intuition behind the enrichment learning algorithm of sparse group-guided RiVIERA-glass model. The learning iterates by alternating between two steps: in step A, we learn the cell-group enrichments using cell group annotations; in step B, we choose to investigate further the cell-type-specific annotations only if the corresponding cell group exhibit prominent enrichment. The output is a sparse interpretable enrichment over the 32 traits investigated in this paper, (c) General framework of the proposed Bayesian fine-mapping RiVIERA-fmap model depicted in probabilistic graphical model (PGM), where the circles are the variables with defined parametric distribution. The outputs are a set of risk loci visualized based on the genetic signals, SNP posteriors, gene tracks, and the top supportive functional annotations.

Using the proposed model, we would like to investigate the following biological questions: (1) where does the functional convergence take place for trait-associated genetic perturbations, as approximated by enrichment profiles of cell-type-specific epigenomic annotations? (2) what groups of traits exhibit similar functional convergence patterns, as measured by correlating enrichment profiles? (3) what are the risk loci supported by both genetic signals and functional enrichment patterns? And (4) what are the specific risk variants within risk loci that exhibit prominent evidence of function? In what follows, we first highlight the novel features of the proposed method. Then we show how such features enable answering these questions directly from real data.

The proposed framework is divided into *two stages* namely genome-wide enrichment learning and locus-specific fine-mapping (**Fig. 1a**). At <u>stage 1</u>, we seek to learn a sparse functional genomic enrichment pattern over all of the tissue-specific annotations with respect to the trait of interest. In other words, we quantify the extent to which SNPs with high genetic signal tend to occur in genomic regions that are functionally active in specific tissues, as indicated by epigenomic annotations. The main challenge here is that many annotations are correlated with each other due to shared basal-level regulatory activities. For instance, in addition to tissue-specific genes, the histone modification H3K4me3 also marks the active promoters ^8,10^ of ubiquitously expressed genes, and it is thus annotated in many cell/tissue types, which are not necessarily related in terms of the underlying tissue-specific biology. Consequently, when jointly modeled in a naïve way, the correlation between different annotations leads to an enrichment pattern that is not easily interpretable. One way to account for the non-tissue-specific enrichment is to include a set of baseline annotations as intercepts in a linear model ^16^. However, although this approach works well when considering one tissue-specific annotation at a time conditioned on the baseline annotations, it becomes inadequate when jointly considering all of the tissue-specific annotations, which nevertheless is necessary for the subsequent risk variant prioritization based on all lines of evidence.

As a novel strategy to these inevitable problems, here we propose a sparse model guided by the enrichments over cell groups pre-defined based on anatomical, hierarchical associations, effectively exploiting the “structure” of the annotation data. Specifically, we iteratively learn the high-level enrichments at the cell-group level and dissect the overrepresentation signal at the cell-type-specific level only within the cell/tissues belonging to highly enriched cell groups (**Fig. 1b**). Importantly, the cell groups are defined based solely on anatomical relations. The annotations in the same cell group are aggregated to form a single annotation, where a SNP is annotated as positive if it overlaps with any of the annotations in the group. Thus, we effectively implement a hierarchical enrichment approach, where signal is quantified first at the organ/tissue level and then propagated at a higher resolution across the relevant cell-types with in the organ/tissue. Our approach is inspired by, but different from, the Group Lasso methods recently developed in statistics ^24 25^. These methods were originally designed to address multi-level categorical variables in a supervised framework where the labels in the response variable are observed, whereas we infer them within the EM framework.

At <u>stage 2</u>, we harness the genome-wide functional enrichments learnt from stage 1 by incorporating them as Bayesian prior into a novel fine-mapping model. As mentioned above, a main challenge of inferring the potential causal variants is the consideration of LD structure. To this end, we need to infer the posterior distributions over *causal configurations* of all of the SNPs in the locus that are potentially correlated with each other via LD. We approach this problem by considering a multivariate normal (MVN) density^17,18,21,26^ as a function of the GWAS summary statistics Z-score with covariance expressed by the reference LD plus the variance contributed by the causal SNPs. However, the brute-force way to model all of the causal configurations is not tractable for modest size locus, because the configurations grow exponentially to the number of SNPs in the locus. In order to circumvent this limitation, we propose an efficient Important Sampling scheme ^21^: we sample configurations based on the posterior probability of each individual SNPs being causal, effectively focusing on the SNPs that are more likely to be causal than the majority of other SNPs in terms of their genetic Z-score (reflected by the likelihood) and functional supports (reflected by the prior).

We validated the proposed approach via simulation (**Supplementary Fig S1-S3**), and in what follows we present our results on the real data.

### 2.2 Functional enrichment profiles of 32 human complex traits reveal relevant non-tissue-specific functional genomic elements

To demonstrate a real-world application, we used the proposed method to investigate 32 human complex traits, for which the full summary statistics are publicly available (**Table 1**). To facilitate downstream analysis, we grouped these 32 traits into 5 groups: Anthropometric, Neuro-degenerative (NeuroDegen), Neuro-psychiatric (NeuroPsych), Heart, Immune, and Metabolic. We preprocessed the GWAS summary statistics by filtering out ambiguous mutations and then imputing them based on 1000 Genome phase 1 version 3 data using ImpG ^27^. In total, we harnessed 272 publicly available and preprocessed functional annotations including 52 baseline annotations and 220 cell-type-specific annotations ^16^.

**Table 1.**
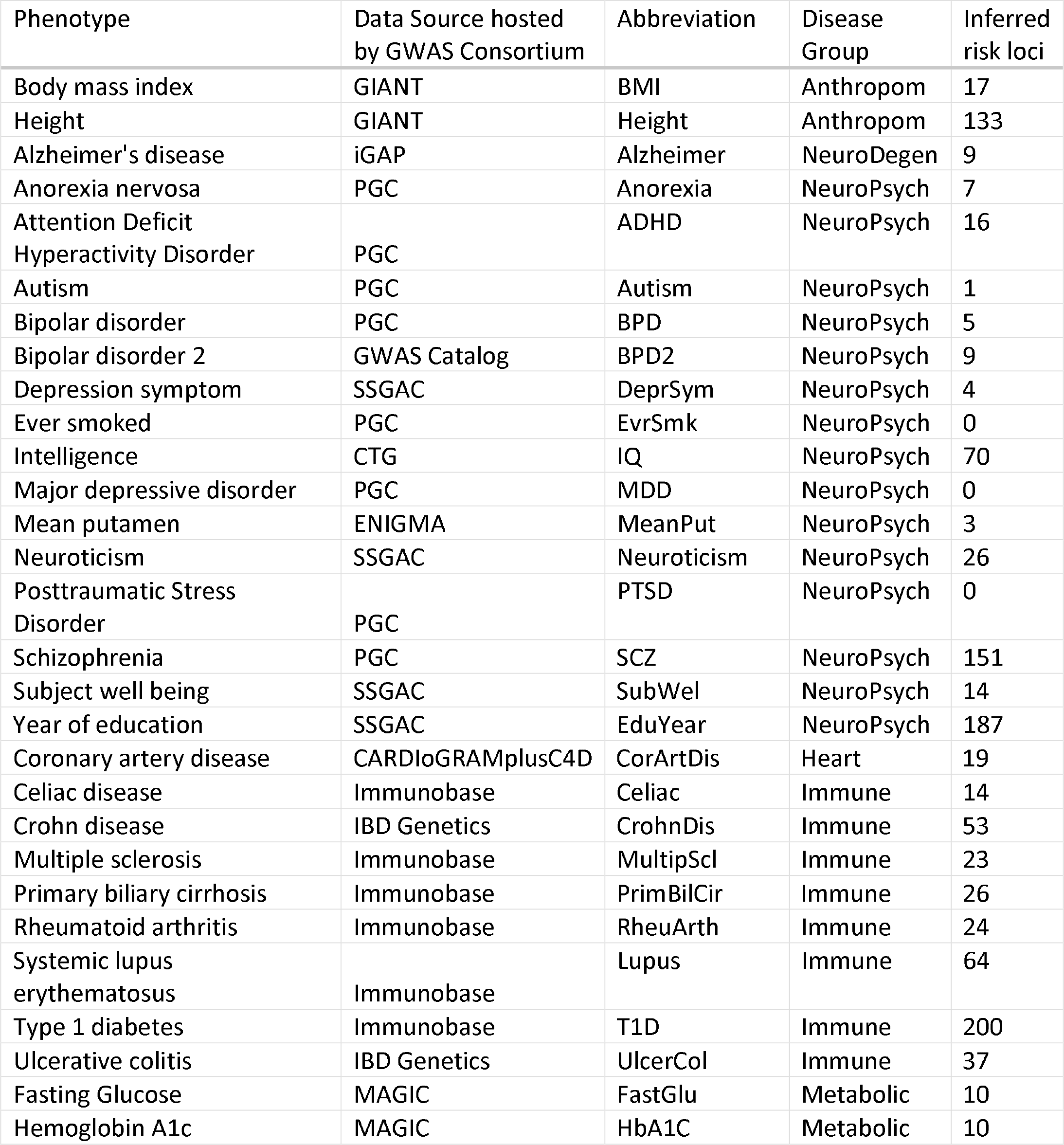
Summary statistics of the 33 GWAS datasets analyzed in this paper

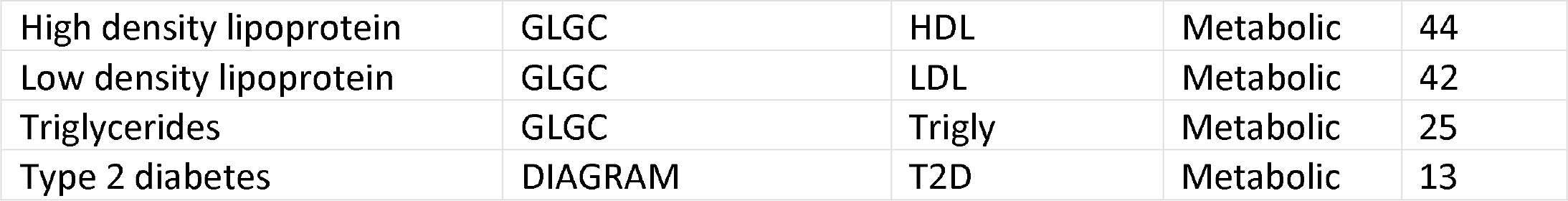

For the genome-wide enrichment analyses, we first performed LD-pruning to recursively remove SNPs that are in high LD with the most significant SNPs to account for inflated p-values due to high LD regions. We then applied our proposed stage-1 model to investigate the functional enrichments of the 32 traits. We first examined what functional categories are enriched for each of the complex traits using only the 52 baseline non-tissue-specific annotations (**Supplementary Fig. S4**; **Supplementary Data 1**). Because sparsity is of less concern with such small number of annotations, we applied a non-sparse variant of our model (i.e., L2-norm over each annotation weights) and assessed significance in terms of the asymptotic Z-score estimates of the enrichment coefficients from the logistic function.

Overall, the most enriched annotations include regions that are transcribed ^28^, conserved in mammals^15^, marked with H3K4me1, overlapping coding segments, and chromatin accessible (DHS). Interestingly, although these 52 baseline annotations are not cell-type-specific, but define a union set of the putative genomic functional space, many traits cluster in a way consistent with the known groups by similarity of enrichment profiles alone. For instance, both immune and brain-related traits aggregate together into clusters. Both immune and metabolic traits tend to be enriched for coding categories. Immune traits also exhibit more prominent enrichments for promoter and enhancers elements (defined by H3K4me3 and H3K4me1/H3K27ac, respectively). In contrast, neuro-psychiatric phenotypes such as human intelligence (IQ), Schizophrenia (SCZ), and Years of Education (EduYear) are strongly enriched for conserved elements and repressed states, and are mostly depleted in coding regions. Thus, the results suggest that overall, genetic variants associated with related traits tend to perturb functionally common sets of genomic elements, and this is evidences even when considering only tissue-agnostic functional annotations.

The strong enrichment for repressed states over many psychiatric traits and their salient contrast to many non-psychiatric traits is intriguing. This state is associated with gene silencing and was originally derived from the repressive histone mark H3K27me3 generated by Polycomb repressive complex 2, which is, for instance, involved in maintenance of pluripotency state during embryonic development ^29,30^. Based on our results, an interesting follow-up study would be to explore whether genetic variants associated with psychiatric traits affect the regulatory roles of this mark converging in subtle alterations in early brain development.

### 2.3 Inferred cell-group enrichments are consistent with the relevant physiology of the complex traits

We then examined the enrichment profiles of the 32 traits for 10 different cell-group annotations. To this end, we aggregated the 220 cell-type-specific annotations into 10 predefined cell-group annotations, namely: adrenal/pancreas, cardiovascular, central nervous system (CNS), connective/bone, gastrointestinal, immune, kidney, liver, skeletal muscle, and others -- according to a previous study^16^. Again, we applied the non-sparse model for this case. We found that our cell-type-specific enrichment estimates are consistent with the underlying biology and the recent literature for most of the traits ^11,16^ (**Fig. 2**; **Supplementary Data 2**). In particular, all of the immune traits exhibit prominent and almost exclusive enrichment for the immune cell-group annotation. Psychiatric traits are predominantly enriched for the CNS annotation with SCZ and EduYear exhibiting the most prominent CNS-enrichments among other related traits. The lipid traits including low/high density lipoprotein (LDL, HDL) and triglycerides (Trigly) are enriched for liver cell group. BMI and Height are mostly enriched for CNS and connective/bone, respectively, whereas fasten glucose (FastGlu) is primarily enriched for adrenal and pancreas. Notably, the genetic signals of ulcerative colitis (UlcerCol) is not only enriched in immune but also in gastrointestinal group, consistent with the underlying pathology ^31^. Interestingly, some of the psychiatric traits exhibit somewhat mixed enrichment signals. In particular, the genetic signals of Alzheimer’s Disease are primarily enriched in immune and liver. Recent literature has suggested both the important roles of innate immunity in Alzheimer^32^ and the metabolic-induced dementia due to liver malfunctions ^33^. The latter may be also related to the enrichment signal we observed for anorexia and autism disorder.

**Figure 2.**
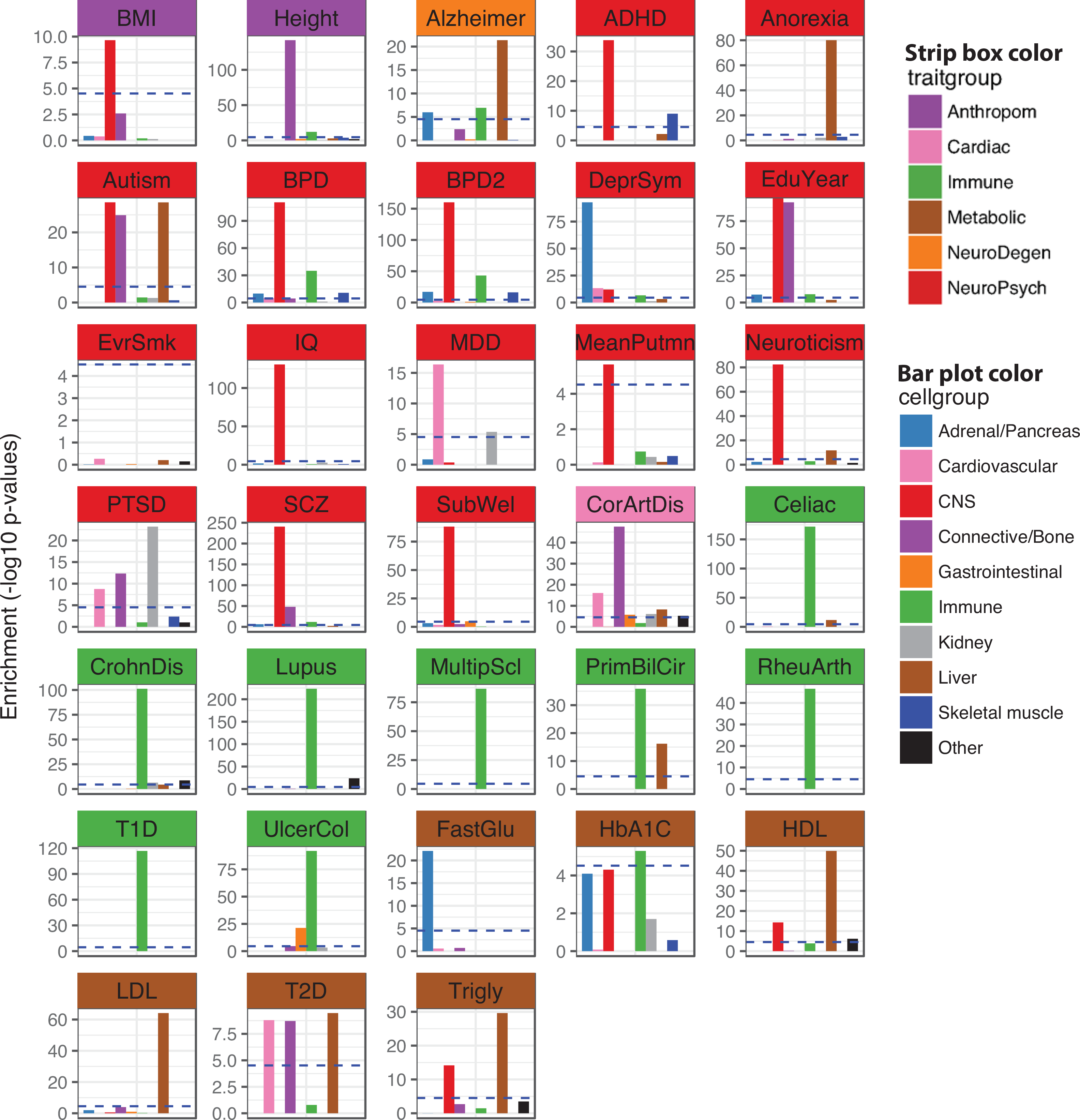
Cell group epigenomic enrichments of 32 distinct traits. Cell-group enrichments were inferred by training the proposed RiVIERA-ridge model over 10 cell-group annotations plus as intercepts the 52 baseline annotations to account for general non-tissue-specific enrichments. The enrichments are displayed as barplot of the −log10 p-values, where the horizontal line indicates p-value < 0.01 after Bonferroni correction for multiple testing (i.e., 330 tests in total).

### 2.4 Sparse cell-group-guided model reveals interpretable enrichment path of tissue-groups

We next proceeded to refine the cell-group enrichments observed above by dissecting the overrepresentation signal at the cell-type-specific level for the highly enriched cell groups. To this end, we used the proposed cell-group-guided sparse model. To examine the effects that different levels of sparsity have on the cell-specific enrichments, we trained our model using values of sparsity ranging from 0.1 to 0.9. We define sparsity level as the fraction of the maximum enrichment score among the 10 cell groups. Thus, a sparsity equal to 0 or 1 will result in the inclusion of either all or no tissue-specific annotations, respectively. To account for general non-tissue-specific enrichments and to facilitate the fine-mapping analysis in stage 2, we also included into our model the 52 baseline annotations (i.e., conservation, coding, transcription factor binding sites, etc.). For the latter we use a different regularization constraint that do not enforce sparsity (i.e., L2-norm).

At each sparsity level, we trained our model until convergence. We then calculated the total enrichment Z-scores over the 220 cell-type-specific annotations by summing the Z-scores estimates over annotations in the same cell group. This resulted in an “enrichment path” (reminiscent of the solution path of Lasso^34^) over the 10 cell-groups as a function of the sparsity thresholds (**Supplementary Fig. S5**). From a Bayesian perspective, perhaps all of the enrichment solutions along the path are valid as they imply different degrees of emphasis over the annotation groups. Nonetheless, we may choose quantitatively the best model based on the penalized log likelihood (**Supplementary Fig. S6**).

It is worth noting that the cell-type-specific enrichments are updated at each iteration only when the corresponding group-level enrichment is above the threshold otherwise they are set to zero. Thus, it is possible that one group of annotations dominates over others at some iterations and later on becomes zero, if their group-level enrichments become less relevant relative to the other groups. Thus, only the annotation groups that persist along the entire iterative learning process will in the end exhibit significant enrichment.

As we can clearly observe (**Supplementary Fig. S5**), the 8 immune traits (from Celiac to UlcerCol panels) display the highest aggregate enrichment scores corresponding to the immune cell group over other annotations consistently across all of the sparsity thresholds tested (i.e., 0.1-0. 9). In contrast, many psychiatric traits such as ADHD, BPD (BPD2), EduYear, IQ, MeanPut, Neuroticism, SCZ, SubWel exhibit predominant enrichment over the CNS annotation group. However, several traits appear to exhibit interesting, combined trends. Bipolar disorder (BPD and BPD2), for instance, exhibits strong enrichments for both the CNS and immune annotation groups up to the highest sparsity threshold, at which only CNS remains. Alzheimer’s disease exhibits strong enrichment in both immune and liver at all sparsity thresholds except for the highest one (i.e., 0.9), where only the enrichment for liver group remains. In contrast, 3 annotation groups namely CNS, connective/bone and liver persist across all thresholds for Autism disorder.

### 2.5 Sparse enrichments identify biologically meaningful cell-type-specific epigenomes indicative of distinct and shared trait biology

We then zoomed into specific epigenomes that exhibit non-zero enrichments for each trait at the most lenient and the most stringent sparsity thresholds that we tested above (i.e., sparsity threshold *λ* = 0.1 and 0.9, respectively) (**Fig. 3a**; **Supplementary Fig.S7**). At both thresholds, the enrichment patterns are striking and biologically meaningful. Although the immune and psychiatric traits are mainly enriched in immune and CNS annotation groups, respectively, individual traits exhibit different enrichments over the cell-specific epigenomes. For instance, at the lenient threshold, UlcerCol is enriched for H3K27ac in both stomach smooth muscle and duodenum mucosa along with highly specific immune cell-types, which are consistent with the disease biology^35,36^. SCZ, IQ, and EduYear are highly enriched for H3K4me1 in fetal brain and H3K27ac in neurophere. Lipid traits namely LDL, HDL, Trigly are enriched in liver-specific epigenomes. Coronary artery disease (CorArtDis) is enriched for adipose nuclei in terms of both H3K27ac and H3Kme1, and it exhibits an enrichment pattern similar to the one of lipid traits, which is consistent with previous findings^37^. At the most stringent threshold (Fig. 3a), the enrichment patterns are more pronounced over the known disease groups and some of the weaker enrichments observed at the lenient thresholds vanish, retaining only the most explanatory epigenomes for each trait. As comparison, we applied a non-sparse variant of the model to the same data and observed much less interpretable enrichment pattern where all of the annotations exhibit non-zero enrichments due to the inter-group correlations (**Supplementary Fig. S8**).

**Figure 3.**
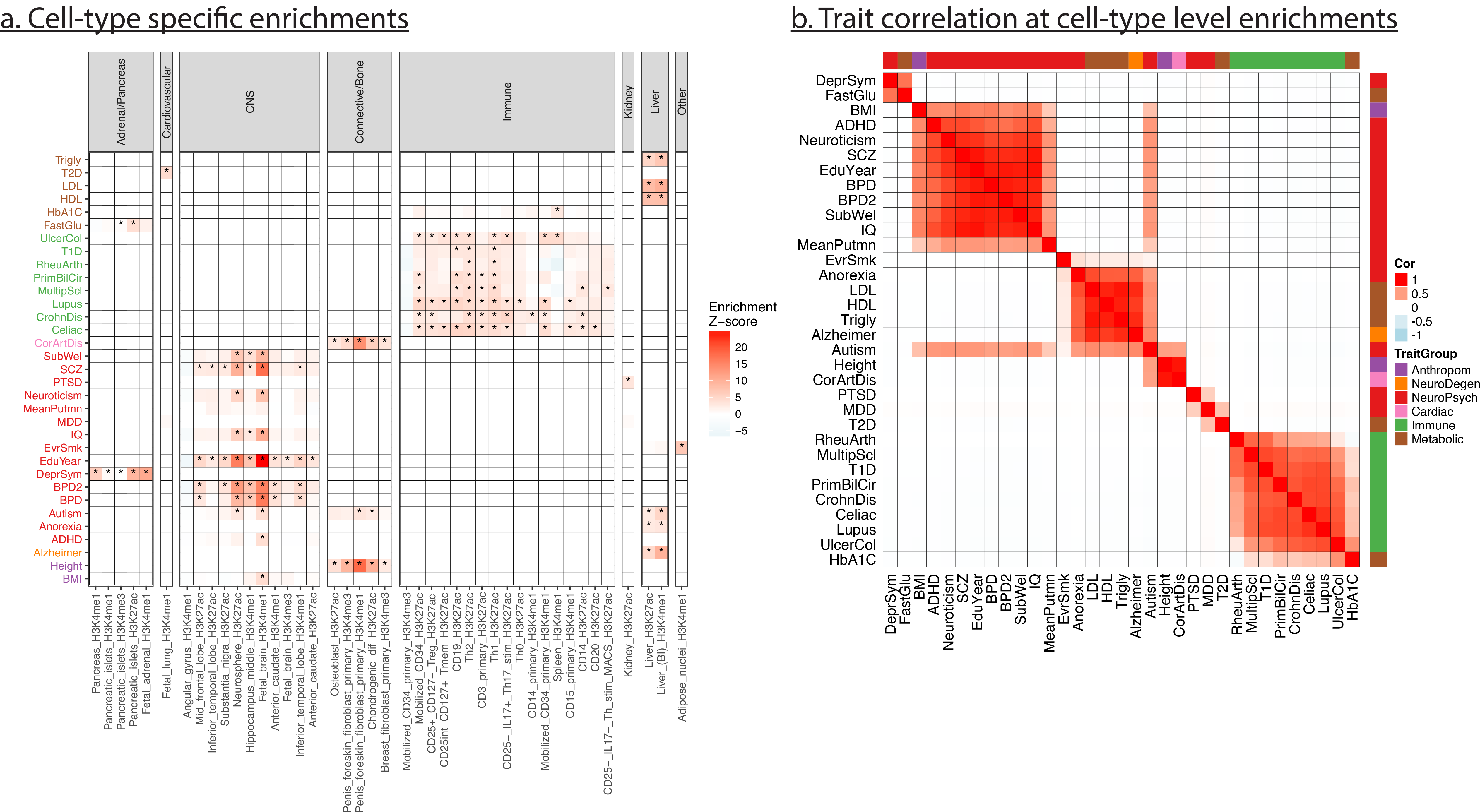
Inferred sparse cell-type-specific epigenomic enrichments. We inferred the enrichments of the 220 cell-type-specific epigenomes for the 32 distinct traits using the proposed group-guided sparse RiVIERA-glass model. The enrichment patterns were displayed as heatmap for the annotations that have non-zero enrichment in at least one of the traits. The colors of the trait text indicate pre-defined disease group to facilitate comparison between traits, (b) Trait-by-trait correlation across the 220 inferred epigenomic enrichments. Traits were ordered based on their correlation. For both plots, the sparsity was set to 0.9 (i.e., the most sparse model among the 9 sparsity tested thresholds). **Supplementary Fig. S7a** illustrates the enrichments of the least sparse model, which gives qualitatively similar results.

We then correlated the traits by their enrichment estimates over the 220 cell-type-specific annotations (**Fig. 3b**; **Supplementary Fig.S7**). At both sparsity thresholds, we observed salient and meaningful disease clustering, largely consistent with the known disease groups. This implies functional co-enrichments of the related traits despite the distinctive GWAS loci associated with many of the related traits we observed in the individual studies. Compared to the low sparsity threshold, the modularity becomes much more prominent at the high sparsity threshold. Intriguingly, Autism exhibits salient co-enrichments with both the lipid traits and some of the psychiatric traits, implying its intricate disease etiology.

We next sought to examine the correlated effects of annotations with respect to each trait. To this regard, we calculated the inverse covariance of the non-zero enrichment coefficients at the lenient threshold (*λ* = 0.9), which is the pairwise partial correlation between annotations whilst controlling for the effect of all of the other annotations. Interestingly, this results in meaningful annotation clusters that are relative to the phenotypes of interest (**Supplementary Fig. S9**; **Supplementary Data 4**). For instance, we observe several annotation modules with respect to human intelligence. In particular, non-tissue-specific annotations such as DHS, DGF, and H3K4me1, and enhancer activities -- all associated with transcriptional activation -- are highly correlated with each other. Tissue-specific annotations related to metabolism such as pancreatic islets and kidney form their own cluster. Notably, the strongest cluster is associated with CNS-specific epigenomes such as H3K4me1 in inferior temporal lobe, hippocampus, anterior caudate and H3K27ac in neurophere, implying a pleiotropic effect where multiple related cell-types are pre-disposed to the disease-causing mutations.

### 2.6 Leveraging functional enrichments to infer risk loci elucidated meaningful trait-associated genes and pathways

To exploit the meaningful genome-wide enrichments learned from the above model, we applied the trained model with the most stringent sparsity (*λ*_*sparsity*_ = 0.9) to score genome-wide SNPs and detect SNPs with high posterior probability of associations (i.e., PPA > 0.6), which exhibit both prominent SNP-level genetic signal and accumulative functional evidence. We define those SNPs as our “lead SNPs”. We chose the most stringent model for the subsequent analyses as it places emphasis on the most important annotations and thus facilitates further interpretations of the prioritized risk variants by the proposed fine-mapping model. Alternatively, one may choose the models that give the highest penalized log likelihood (**Supplementary Fig. S6**). We then constructed risk loci by including SNPs within 100-kb distance from the lead SNPs, merging overlapping loci. As a result, we identified 1256 risk loci in total for 29 out of the 32 traits (**Table 1**). Notably, many of the risk loci do not harbor genome-wide significant SNPs (i.e., p-value < 5e-8) but all of them exhibit sub-threshold significance level (p-value < le-5) (**Supplementary Data 5**).

We then examined how these risk loci cluster diseases. To this end, we obtained 1703 approximately independent LD block based on 1000 Genome European population using LDetect from recent study^38^ and represented the genetic signal of each LD block by the highest PPA from the overlapping risk loci such that each trait has 1703 PPA values representing a distilled version of the genome-wide associations for them. We observed that the phenotypic cluster based on PPA is largely consistent with the known disease group (**Supplementary Fig. S10**). This implies some degree of co-localization of risk loci for similar phenotypes. Notably, some traits such as T1D, SCZ, EduYear, and Height are more polygenic than other traits.

We then investigated the genomic location of the risk loci and annotated them based on the nearest protein-coding genes from the lead SNPs harbored in the loci (**Supplementary Fig. S11**; **Supplementary Data 5**). Many genes are shared among related traits (**Fig. 4**). These include genes that are nearby sub-threshold risk loci. These shared genes are closely related with the biology of the traits. For instance, *Interleukin 10* (*IL10*), which we found to be associated with CrohnDis, UlcerCol, and T1D, is an anti-inflammatory cytokine. *Myocyte Enhancer Factor 2C* (*MEF2C*) is known to play a role in maintaining the differentiated state of muscle cells ^39^. However, mutations and deletions at this locus have been associated with severe mental retardation, stereotypic movements, epilepsy, and cerebral malformation. Remarkably, we found *MEF2C* associated with 6 traits: Neuroticism, EduYear, IQ, SCZ, ADHD, and Height. We will further investigate the *IL10* and *MEF2C loci* in the next section of the fine-mapping analysis.

**Figure 4.**
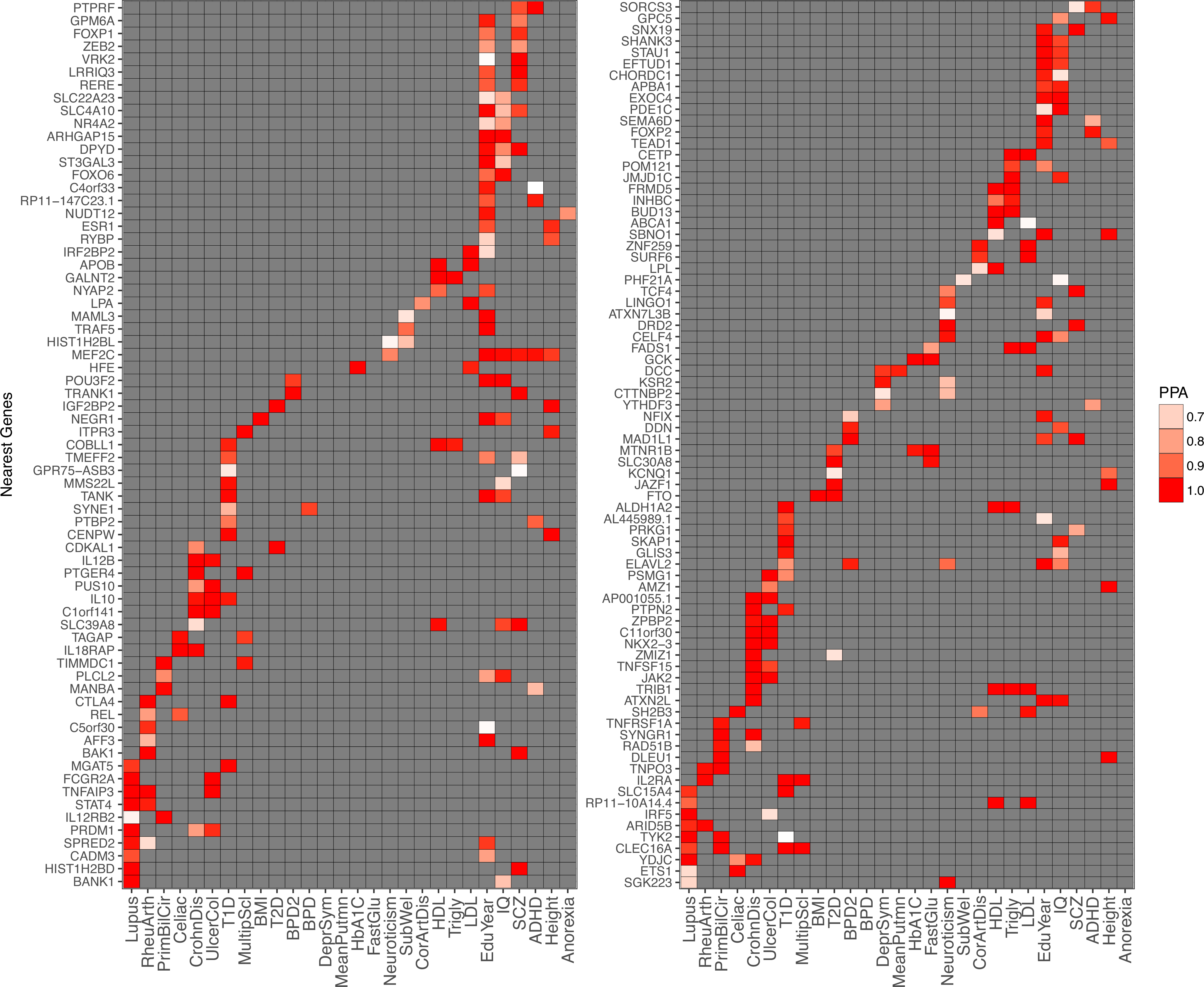
Risk genes associated with two or more traits. We annotate the loci by the nearest genes and obtained risk genes that are common in at least two traits. Genes nearby risk loci are enriched for biologically meaningful pathways and gene ontology terms for select traits.

In order to further test whether the identified associated genes tend to converge into biologically meaningful functional classes or known pathways, we performed gene set enrichment analysis on a per trait basis, using the genes nearby the risk loci. Interestingly, overall, we found that the biological processes (BP) and pathways detected as significant are consistent with disease biology (**Fig. 5** and **Supplementary Fig. S12**). For example, the three most enriched BP for (1) Schizophrenia, (2) Years of education, and (3) Multiple Sclerosis are, respectively: (1) modulation_of_synaptic_transmission (p-value < 1.9e-4), regulation_of_synaptic_potentiation (p-value < 3e-3), and neuron_differentiation (p-value < 3.8e-3); (2) neuron_projection_morphogenesis (p-value < 1.7e-5), neuron_differentiation (p-value < 1.7e-5), neuron_projection_development (p-value < 5.6e-5); and (3) regulation_of_inflammatory_response (p-value < 3.4e-3), defense_response (p-value < 6.6e-3), and response_to_wounding (p-value < 6.6e-3). These results are consistent with the known developmental/psychiatric, cognitive, and immune nature of the diseases. We include enrichment results for all traits in **Supplementary Data 6** and **7**. Furthermore, under the hypothesis that the identified genes are likely to have a functional influence under different types of perturbations, we analyzed whether the latter also presents functional constraint as evidenced by depletion of deleterious of coding variation within humans. Using recently published scores for loss-of-function (LoF)-intolerance estimated from human exome data ^40^, we found that identified genes are under significantly higher constraint compared to expectation (p-value < 2.2e-16; **Fig 5b**). Further, we classified the identified genes in two groups according to whether they were found to be associated with 2 or more traits (pleiotropic) or only one (non-pleiotropic). As expected, both pleiotropic (p-value < 3.6e-6) and non-pleiotropic (p-value < 4.5e-4) genes have significantly higher constraint scores than expectations. Interestingly, however, pleiotropic genes have a more extreme deviation (**Fig 5b**). Thus, the identified genes tend to be more depleted of loss-of-function coding variation relative to background expectation, and the degree of constraint is greater in those associated with multiple traits. These results further support the potential functional relevance of the identified genes and the effects of their disruption by mutation.

**Figure 5.**
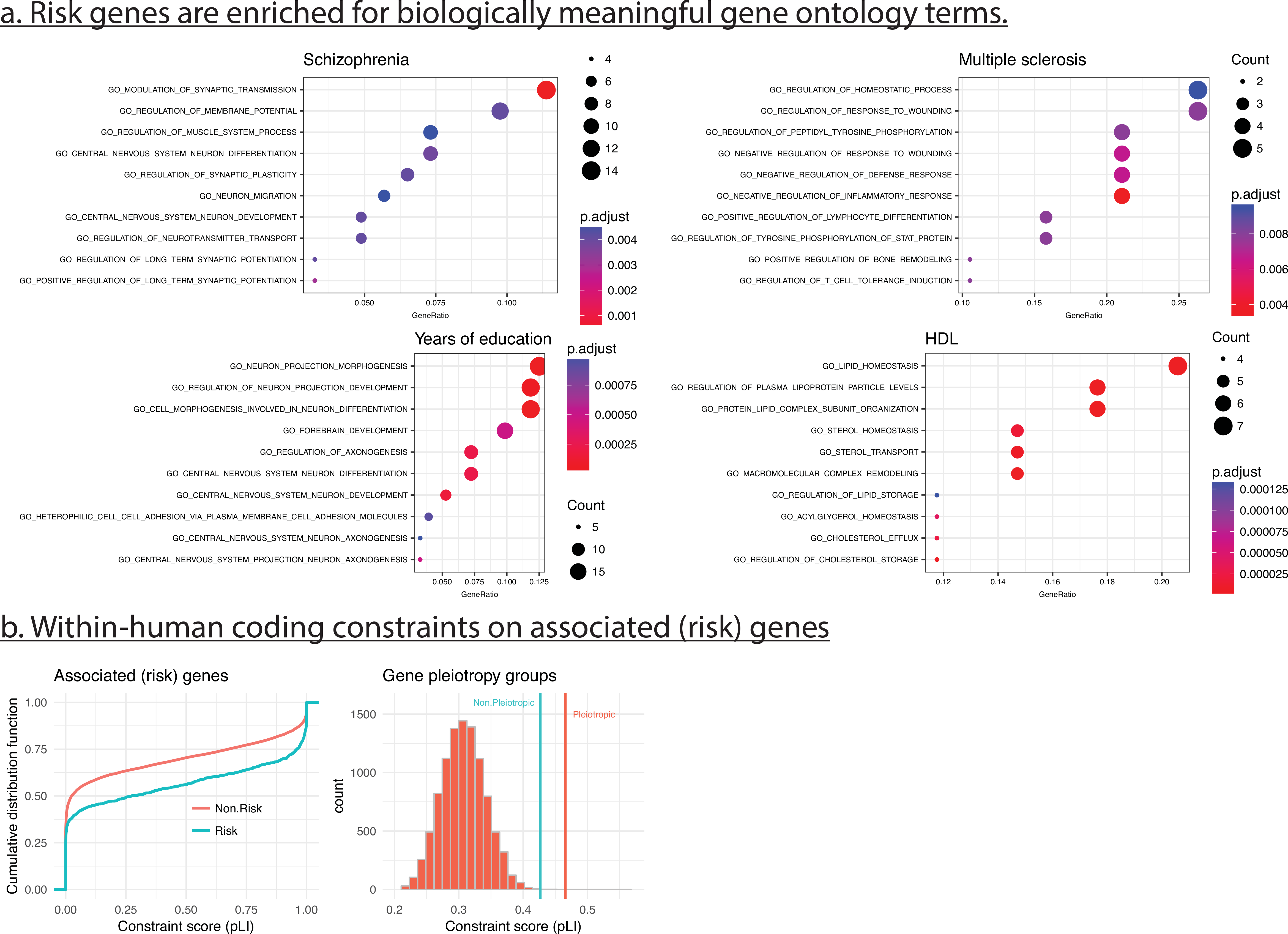
Functional enrichments of the risk genes. (a) Genes nearby risk loci are enriched for biologically meaningful pathways and gene ontology terms for select traits. Gene set enrichment analysis shows significant over-representation of biological processes (BP) and pathways consistent with disease biology, (b) Within-human coding constraints on associated (risk) genes. Genes nearby risk loci are under significantly higher constraint than background (non risk) genes, as estimated exome-based loss-of-function (LoF)-intolerance scores. Genes associated with more than one trait (Pleiotropic) have a more extreme deviation than risk genes associated with only one trait (Non-pleiotropic). All identified risk genes as a group have significantly higher constraint scores than expectations.

Overall, results in this section indicate that our integrative approach recovers biologically meaningful genetic associations that converge to candidate genes with likely functional and tissue-specific consequences under disruption.

### 2.7 Integrative fine-mapping prioritizes risk variants based on tissue-specific evidence of activity

To infer causal variants within the risk loci, we further refined the loci by including only the SNPs within modest LD of (|*r*| > 0.2) with the lead SNPs (PPA > 0.6) for each locus as the remaining SNPs are unlikely to underlie the association. Note that we retained SNPs with absolute Pearson correlation greater than 0.2 with the lead SNPs to account for opposite effect and to better model the corresponding z-scores with a multivariate normal distribution.

We then applied our proposed fine-mapping model (RiVIERA-fmap) to infer the marginal posterior distribution of the SNPs being causal within each of the risk loci as well as the locus-specific enrichments of the reference annotations (**Supplementary Data 5**; **Table 2**). The latter differ from the genome-wide estimates due to the more confined set of SNPs and the result of disentangling causal SNPs from LD-linked SNPs. The inference here is challenging because the number of SNPs is much lower in the risk loci relative to the full genome, and the non-causal SNPs are often linked with causal SNPs via LD. Therefore, we sought to approach such uncertainty by using a novel Bayesian framework (**Fig. 1c**), where the posterior distribution of causal SNPs and enrichment coefficients are approximated by sampling.

**Table 2.**
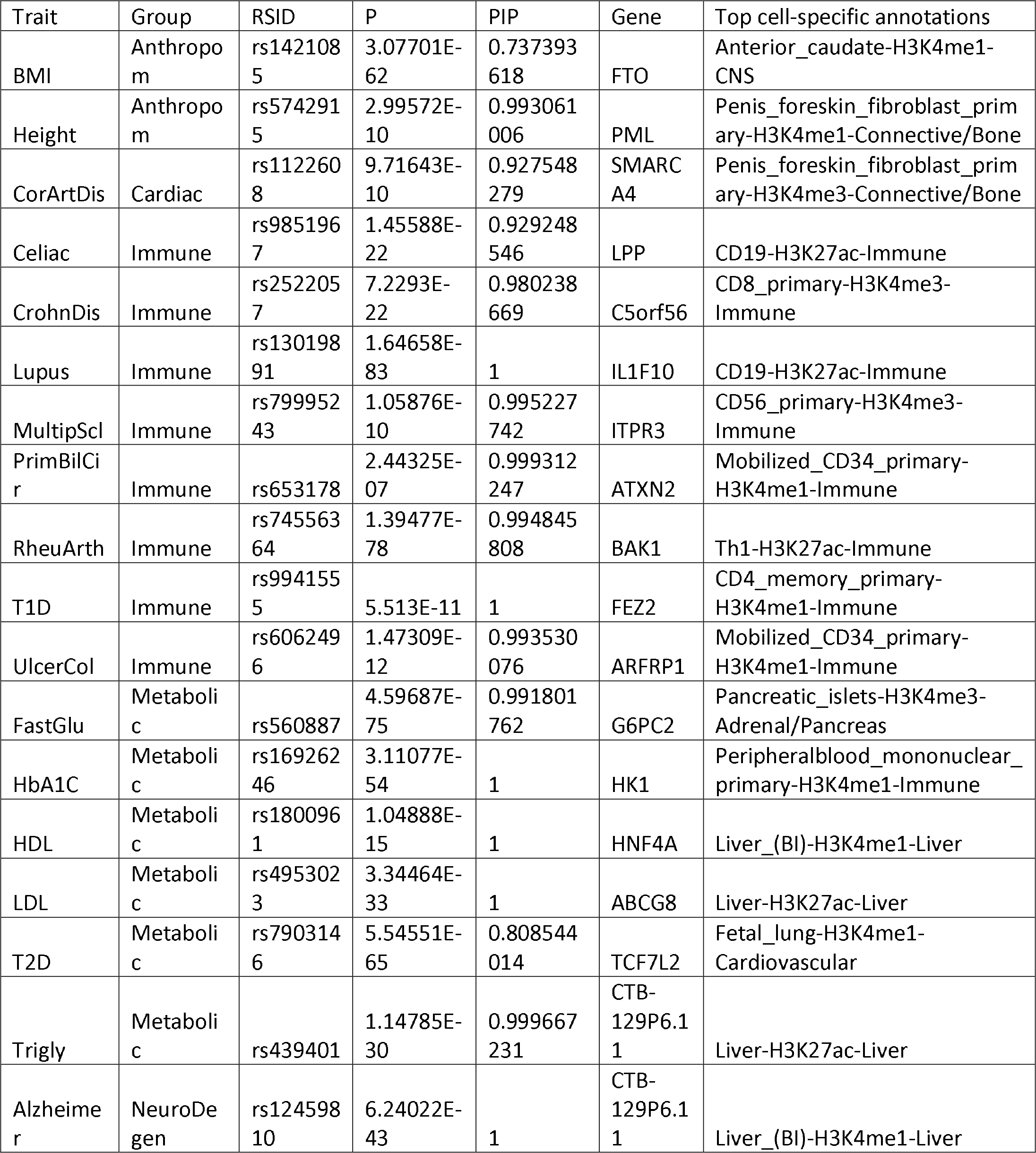
fine-mapped top loci for each trait

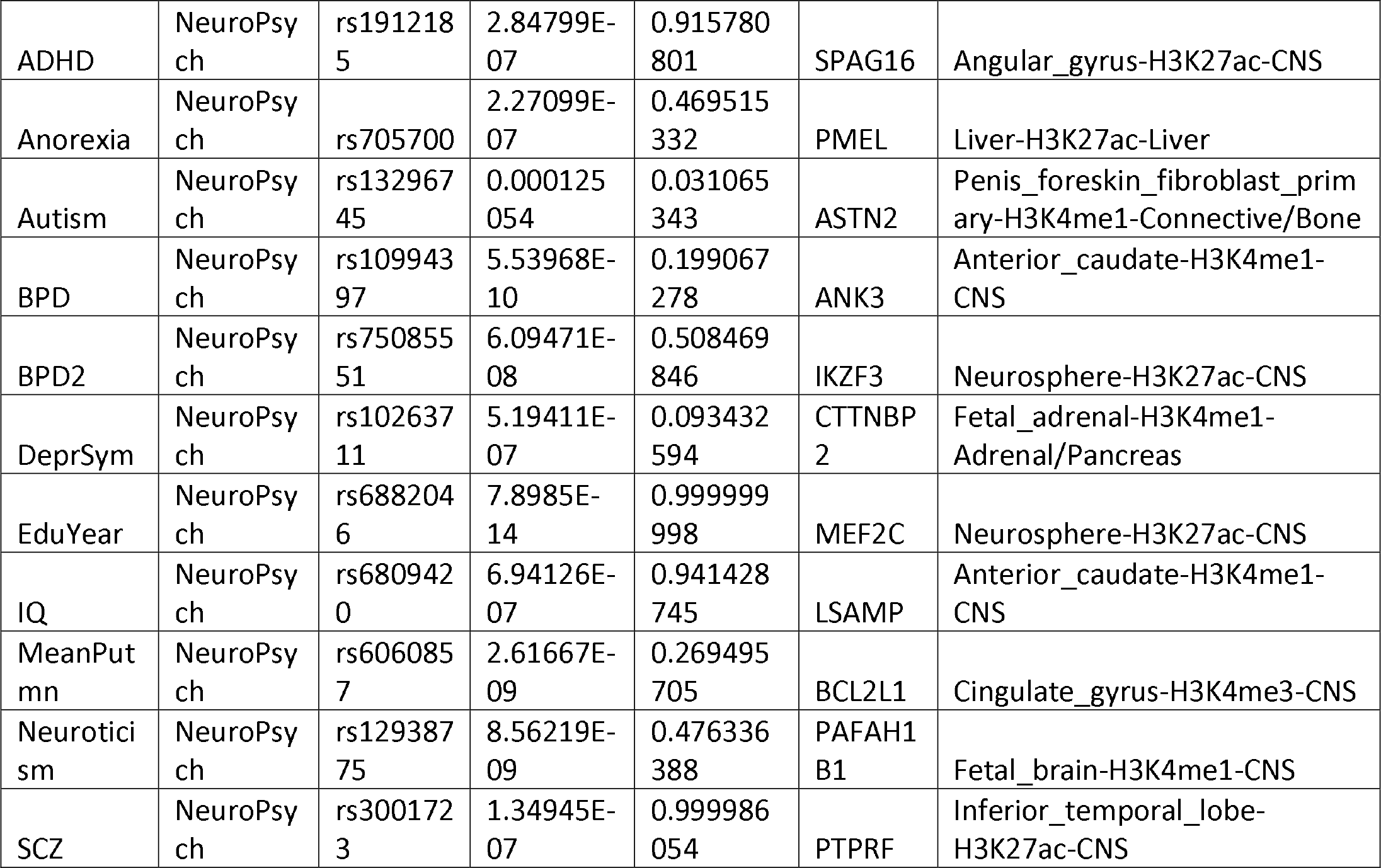

Specifically, we modelled the prior of locus-specific enrichment coefficients with a multivariate normal distribution, where the mean (i.e., **Fig. 3**) and annotation covariance (i.e., **Supplementary Fig. S9**) were fixed to the ones learned during the global genome-wide estimates inferred (stage 1 of the framework) above. Annotations with zero enrichment coefficients in the above genome-wide sparse model are not considered in the fine-mapping model. We performed Hamiltonian Monte Carlo (HMC) ^41^ to sample from the corresponding posterior distribution to approximate the enrichment distribution. We ran our model for 5000 MCMC iterations. At each iteration, we sampled 100 configurations per locus by Important Sampling and calculated the posterior inclusion probability (PIP) of SNPs being causal by marginalizing the posteriors over the sampled and weighted causal configurations whenever a new set of enrichment coefficients are sampled at the HMC step. The final PIP values are averaged to obtain the Bayesian estimates of the SNP causal status.

We first assessed the functional implication of the SNPs by testing their overlap with independent evidence suggestive of function. First, we calculated the significance of overlap of the prioritized SNPs with eQTL SNPs from GTEx (version 6) whole-blood samples. To account for LD in the eQTL data, we retained only the SNPs in both eQTL and digital genomic footprint (DGF) sites at 6-bp resolution derived from DNaseI data ^42^. We observed that the prioritized SNPs taking the top 100 to 500 SNPs based on estimated PIP posteriors are significantly more enriched for eQTL compared to prioritized SNPs based solely on the GWAS genetic signals (− log *p*) from GWAS alone (Wilcox signed-rank test p < 0.007, **Fig. 6a**). Next, we examined whether the prioritized SNPs are relatively more evolutionarily conserved. We used the PhastCons46Way conservation score based alignments across 46 species obtained from UCSC genome browser, and calculated for the top rank SNPs the average conservation. Indeed, our prioritized SNPs exhibit significantly higher conservation compared to the SNPs prioritized by the GWAS p-values (Wilcox signed-rank test p < 1.72e−6, **Fig. 6b**). These results suggest that the additional information provided by tissue-specific evidence of regulatory activity (i.e., epigenomic annotations), indeed seems to help finely localize variants more likely to have a functional impact. This observation thus highlights the utility of having a method able to automatically integrate genetic and epigenetic evidence, while at the same time pointing to likely tissues-of action.

**Figure 6.**
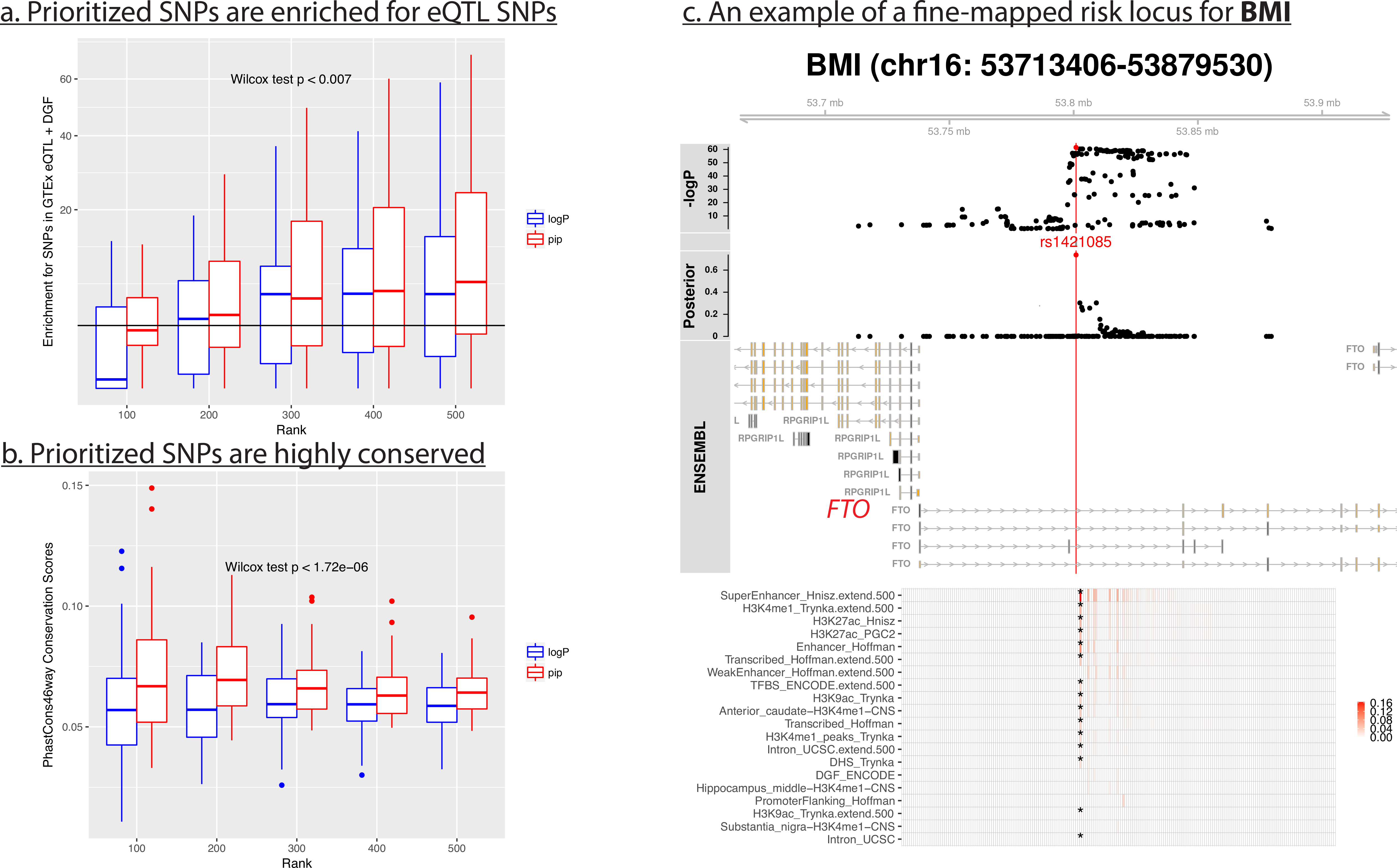
Prioritized SNPs are enriched for functional elements in the risk loci. We performed fine-mapping using the proposed Bayesian RiVIERA-fmap model for each trait and examined the prioritized SNPs based on the posterior inclusion probability (PIP) for their functional evidence, (a) Prioritized SNPs are enriched for eQTL SNPs. We took the top 100 to 500 SNPs based on either the GWAS −logP or PIP. At each ranking, we assessed the hypergeometric enrichment of the SNPs for eQTL SNPs obtained from GTEx whole blood samples. The Wilcox rank-sum onesided test was performed by comparing the enrichments scores from PIP-ranked SNPs with the those from SNPs ranked by −logP. (b) Prioritized SNPs are highly conserved. Similar to (a), we compared the average conservation score from PhastCon46way for the top ranked SNPs by either GWAS −logP or PIP. (c) Fine-mapping example for the *FTO* locus. The tracks from top to bottom display respectively the −logP genetic signals, top SNP within each locus, posterior of the SNPs, Ensembl genes, and the top 20 annotations weighted by the PIP of the SNPs in the loci and the averaged enrichments inferred by the model across all loci.

### 2.8 Examples of fine-mapped loci

We then zoomed into the specific loci to see what SNPs are attributable to the observed locus-specific functional enrichments (**Table 2**). To this end, we visualized the inference results on a locus-by-locus basis and identified many interesting high-risk non-coding SNPs that potentially disrupt the regulation of nearby genes. Interestingly, many of the genes are also highly expressed in the relevant tissues, consistent with the epigenomic enrichment. For the sake of concreteness, we are going to discuss only five loci, considering traits from the five predefined phenotypic groups.

As the first example, we zoomed into the *FTO* locus, which based on the present analysis is associated with both BMI and T2D (**Fig. 4**). *FTO* is known to be associated with obesity-related traits ^43 44^. The locus is within a tight LD block, including many SNPs that exhibit high significant p-values (**Fig. 6c**). Nonetheless, by modeling the effect of LD and leveraging the genomic and epigenomic enrichments, we prioritized a SNP (rs1421085) that is supported by many lines of functional genomics evidence, including overlap with super enhancer, H3K27ac, H3K4me1 activity in brain anterior caudate region. Moreover, the SNP is also correlated with the *FTO* gene expression in skeletal muscle samples from GTEx, thereby providing further evidence to the specificity of its functional potential. Importantly, the very same SNP has been recently experimentally validated by our group ^44^. In particular, the T-to-C single-nucleotide mutation at rs1421085 disrupts a conserved motif for the ARID5B repressor, which leads to derepression of a pre-adipocyte enhancer and a two-fold increase of *IRX3* and *IRX5* expression during early adipocyte differentiation ^44^. This fact, strongly demonstrate the utility of the proposed framework, as it learnt in a completely unbiased and automatic way a non-coding genetic perturbation that has previously shown to mechanistically lead to a cellular phenotype directly impacting obesity. Thus, the identified variants constitute testable hypotheses about the mechanisms underlying the genetic association.

As the second example, we focus on a risk locus (chr1: 25661137-25860915) harboring four comparable SNPs, which exhibit strong evidence of enhancer (H3K4me1/H3K27ac) activity in liver tissues (**Supplementary Fig. S14a**). The nearest gene to these variants is *TMEM57*, which encodes a membrane protein with poorly characterized function ^45^. Interestingly, *TMEM57* is also the proposed target gene of the top SNP (rs10903129), prioritized based on GTEx eQTL analysis in several relevant tissues including aorta and tibial artery tissues. Notably, LDL cholesterol level is one of the main causes of cardiovascular disease ^45 46^ but the potential role of *TMEM57* in both traits remains to be elucidated. However, we do not rule out the possibilities that genes other than the nearest might be affected by the same SNP. As proposed by recent transcriptome-wide association studies, there were cases that the nearest gene is not necessarily the target gene of the variants ^47^. For instance, note that *LDLRAP1* (Low Density Lipoprotein Receptor Adaptor Protein 1) is also within close vicinity of the same locus and thus may also be affected by the mutation although we could not find corroborating GTEx eQTL results. We also examined the inferred posterior distribution of the functional enrichments of both the baseline and tissue-specific annotations and found that the most enriched annotations for LDL are extended coding regions and H3K27ac activity in liver (above 90% Bayesian credible interval) (**Supplementary Fig. S13**).

As the third example, we illustrate *IL10* locus, which exhibits a pleiotropic association with three related traits (CrohnDis, UlcerCol, and T1D) (**Supplementary Fig. S14b**). Four SNPs in this locus exhibit not only high genetic signals but also prominent epigenomic signatures in immune cells, implying functional relevance to the immune disorders. As the fourth example, we examined the *NBEAL1* locus associated with CorArtDis. *NBEAL1* is known to be associated with myocardial infarction gene ^48,49^. Because of the extensive LD (**Supplementary Fig. S14c**), identifying the causal variants based only on the GWAS genetic signals is difficult. However, when combining with various functional genomic and epigenomic reference annotations as well as modelling the z-score distribution as a function of the LD reference panel, we identified only one variant (rs2351524) with high posterior probability (PIP > 0.6). Remarkably, the SNP is also associated with the gene expression of *NBEAL1* based on GTEx data from coronary and aorta artery samples, as well as the nearby gene *CARF* in atrial appendage heart. This observation supports a likely disruptive role of the prioritized variant in the regulation of gene expression specific to disease relevant tissues.

Finally, we highlight the gene locus *MEF2C*, which interestingly was found to be associated with four different cognitive/psychiatric traits (EduYear, IQ, SCZ, ADHD) as well as human height (**Fig. 4**). This finding demonstrates the benefit of analyzing and comparing multiple traits in parallel, which enables the discovery of promising pleiotropic loci. *MEF2C* codes for myocyte-specific enhancer factor 2C, a transcription activator that binds specifically to the MEF2 element present in the regulatory regions of many muscle-specific genes ^39^. However, it is known to play an important role in hippocampal-dependent learning and memory, by suppressing the number of excitatory synapses and thus regulating basal and evoked synaptic transmission ^50^. It is also crucial for normal neuronal development, distribution, and electrical activity in the neocortex ^51^. We prioritized a top SNP (rs41352752) in this locus, which, for example, in the IQ-associated locus exhibits much stronger functional evidence or brain regulatory activity, such as super enhancer, H3K27ac in inferior temporal and middle frontal lobe, relative to other SNPs with comparable genetic signal in the same locus (**Supplementary Fig. S14d**). Again, this example demonstrates how our integrative modeling framework is able to provide functionally supported hypothesis for experimental follow-up studies.

## 3 Discussion

Dissecting molecular mechanisms underlying genetic association with complex traits, ultimately mapping genotypes to phenotypes, is becoming more feasible thanks to the recent availability of large-scale functional genomics data ^8,52–54^. One natural approach is to incorporate various genome-wide reference annotations in the form of prior evidence within a Bayesian framework to infer the functional variants that underlie the genetic signals ^11,19,20,23,55,56^. However, given hundreds of collinear annotations, it is challenging to identify and interpret those annotations that are more specifically enriched with genetic signal. This limitation reflects the mixed success while trying to do so ^16,19,56,57^. Moreover, it is often difficult to harmonize the identification of causal variants (i.e., fine-mapping), which are restricted to only a set of loci ^21,23,58^, with the specific identification of genome-wide relevant annotations ^16^. Here we propose a novel efficient framework to address that challenge.

Our main contributions are the following. <u>First</u>, using the proposed efficient genome-wide model, we demonstrate that overall the traits we investigated exhibit meaningful and statistically significant enrichments for tissue/group-specific epigenomes when modeled together. <u>Second</u>, using a novel sparse group-guided model, we identified significant cell-type-specific epigenomes that are highly relevant to the traits of interest, thereby suggesting the target cell types of action for follow-up experimental investigation. Moreover, we showed that the majority of the traits aggregate in clusters based on the epigenomic enrichment profiles, and that the uncovered clusters are consistent with known trait/disease groups. Because many of these traits do not share the same risk loci, our results imply a model in which genetic perturbations associated with complex traits converge into downstream regulatory mechanisms that are shared across related tissues. Furthermore, we demonstrate a novel way to detect modules of functional annotations with respect to a target trait, by exploiting the covariance of the enrichments directly from the trained model. One caveat of this approach is the assumption of traits being measured from non-overlapping samples, as we assume that z-scores are independent across traits, which is not necessarily the case. While independence seems to hold for most traits that we investigated, we cannot eliminate the possibility that common disease clusters are indirectly related to overlapping samples between some traits.

<u>Third</u>, we harness the meaningful enrichments that we learned by incorporating them as priors in a novel Bayesian fine-mapping model. As a result, the prioritized variants exhibit not only strong genetic signals but also strong evidence of tissue specific regulatory activities. This is especially important for many cases where the causal variants are in a strong LD with other variants. For instance, we are able to recapitulate the causal variant in *FTO* locus that was previously validated in a mouse model to have causal effect on obesity by analyzing the publicly available BMI GWAS summary statistics. In addition, there are many variants that we prioritized and appear to be as promising as the validated SNP based on multiple lines of supporting evidence, from both genetic and epigenomic data, thereby providing a valuable repertoire of hypotheses for the research community interested in experimentally dissecting GWAS loci.

Additionally, RiVIERA is implemented as an R package so that it can be easily to apply to other datasets. As inputs to the software, users need to supply RiVIERA with summary statistics, reference annotations, and LD information to estimate genome-wide functional enrichment estimates, infer risk loci, fine-map risk variants all with the same software package. As for the computational resource needed to run the full analysis described in the paper, the genome-wide model will require at least 32 GB RAM to load 1-5 million SNPs and hundreds of annotations in memory, whereas the fine-mapping model require much less memory for the refined set of risk loci. Both models typically finish in a few hours on a single-core modern CPU.

As future works, the current framework can be extended in many ways. First, causal genes could be inferred beyond individual variants by jointly modeling both the GWAS data and eQTL data taking into account not only the SNP-level annotations but also gene-level network information such as signalling pathways or protein-protein interactions. Second, for traits that are associated with the same pleiotropic loci, we may provide the users an option to infer the joint posterior of SNP associations with multiple traits ^21^. Third, the current model can also be easily adapted to model trans-ethnic GWAS using separate LD matrices, as effectively demonstrated by the trans-ethnic version of the PAINTOR model^22^. Fourth, instead of using the linear logistic prior model, we will explore other models that take into account the spatial information of the genomic sequence as well as local epigenomic context around each SNP. Fifth, the group-guided sparse model can be further improved by imposing richer structure such as a hierarchical tree structure over the annotations with increasing cell-type-specificity going from the root to the leaf nodes. Finally, to facilitate dissemination of our fine-mapping results, we will consider the possibility of having an on-line portal for users to explore the model inference in an interactive way. Thus, the present work provides a useful building block for more complex modeling tasks and user-friendly applications for GWAS dissection.

All together, we consider that the proposed framework provides a valuable demonstration of the power gained by integrating functional epigenomic data from ENCODE and Roadmap Epigenomics Consortia, population genetic data in terms of LD from 1000 Genome Project, tissue-specific genotype-expression information, and genetic signals from large GWAS consortia in order to dissect genetic signal in terms of specific molecular mechanisms, hereby going from an agnostic data-driven approach to a refined set of specific biological hypotheses that constitute experimentally actionable items in dissecting functional genetics.

## 4 Methods

### 4.1 Learning genome-wide epigenomic enrichments

To detect genome-wide functional genomic and cell-group-specific enrichments, we propose an efficient mixture model learning approach with expectation-maximization updates. In particular, we developed two models namely (1) RiVIERA-ridge and (2) RiVIERA-glass. The first model RiVIERA-ridge does not impose sparsity and uses L1-norm constraint on the enrichment coefficients, which works well on small number of approximately independent annotations, whereas the second RiVIERA-glass model builds upon RiVIERA-ridge and exploits group-level sparsity, which confers much more interpretable enrichment estimates over hundreds of the cell-type-specific annotations. Below we describe the main components of the two models. Interested readers can refer to extended details in **Supplementary Note**.

#### 4.1.1 Non-sparse RiVIERA-ridge model

We model the p-values of the SNPs using a two-component Beta mixture model, which is similar to some of the previous works ^20,56,59^. The null and signal component correspond to whether the SNPs are in or outside of a risk locus, which is reflected by the high and low p-values, respectively. Here we assume that the SNP signals ***x*** are conditionally independent given their latent indicators **z** (for being in the risk loci or not). Specifically, suppose there are N SNPs, the *expected complete log likelihood* is defined as follows:

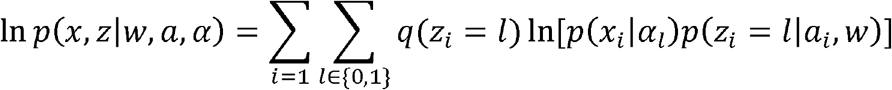

where

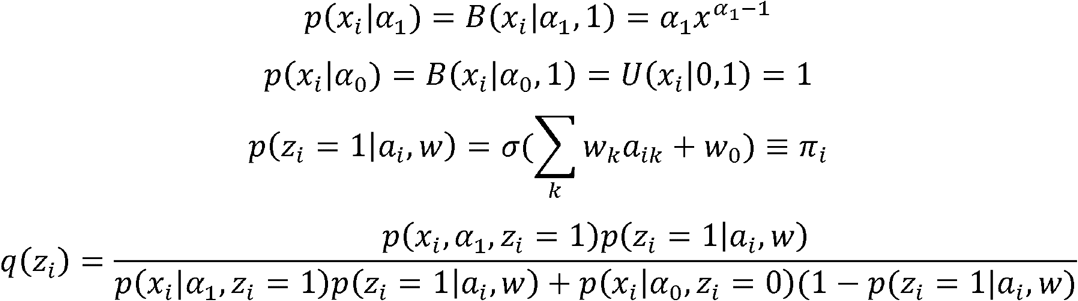

where *σ*(*y*) = 1/(1 + exp(−*y*)) and *α*_0_ and *α*_0_ are the null and causal mean p-values for the non-risk and risk locus, respectively. We estimate *α*_1_ from the data and fix *α*_0_ to 1, which is equivalent to uniform distribution. The parameters **w** = [*w*_0_, *w*_l_,…,*w*_*k*_]′ are the intercept and enrichment coefficients for the K annotations. And *q*(*z*_*i*_) = *p*(*z*_*i*_|*x*_*i*_, *α*, *a*, *w*) is the posterior probabilities of association (PPA)^60^.

By applying the Bayes rule, the posterior probability of the model parameters is expressed as follows:

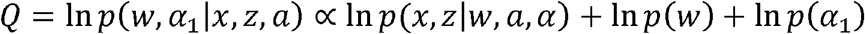

Here we assume In *p*(*w*)=∑_*k*_ ln*N*(*w*_*k*_|0,1) and In*p*(*w*_0_)=In**N*(*w*_0_|*μ*_0_*, 1) and *p*(*α*_1_) follows a uniform prior and its value does not affect the posterior as long as it is greater than zero. *μ*_0_ =log(*π*_0_) − log(l − *π*_0_), where *π*_0_ = 0.001, implying *apriori* the probability of a SNP being a risk-associated variant is 0.001.

Our goal is to optimize the objective function *Q* with respect to the enrichment parameters **w**. Notably, the optimization procedure requires the evaluation of *q*(*Z*_*i*_), which in turns depend on the model parameters. Thus, we use EM algorithm to update these parameters:

1. Initialize *α*_1_ =0.1, *π*_0_ = 0.001, *μ*0 = *w*_0_ = ln(*π*_0_) − ln(1 − *π*_0_);
2. E-step: update *q*(**z**);
3. M-step Update *α*_1_ and **w** by Newton-Raphson update;
4. Repeat step 2 and 3 until convergence.

Because both the E-step and M-step maximize the objective function, the algorithm guarantees convergence.

### 4.2 Significance testing for functional enrichments

To test for the significance of function enrichment for annotation *k*, we compute the corresponding z-score from the enrichment coefficients 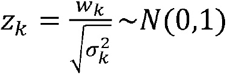, where 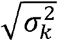 is the *k*^*th*^ diagonal element of the covariance matrix *∑*_*w*_, which is the inverse of the negative Hessian matrix estimated using the converged model.

### 4.3 Group-sparse model to learn cell-type-specific enrichments over hundreds of epigenomic annotations

To learn a cell-type-specific enrichment model, we proposed a group-guided update algorithm. The objective function of the model parameters specified above stays the same except for the change of the prior constraint over **w** from univariate Gaussian (or L2-norm) to a constraint function that leverages group-level information over the annotations (i.e., L1/L2-norm):

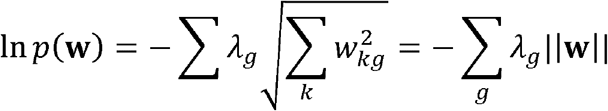

which imposes a L1-norm over each annotation group and a L2-norm over annotations within each group. The gradients of the objective function with respect to the annotation enrichment weights are defined everywhere except when 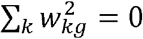. Based on KKT condition, the optimal solution for group *g* is zero if 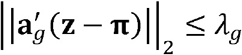 ^24,25^.

However, the standard Group Lasso approach does not work well for our problem in practice due to the highly unbalanced number of annotations in each cell group and perhaps nonstandardized binary annotation variables in order to facilitate interpretations. For instance, there are 67 immune epigenomic annotations, 34 CNS epigenomic annotations, and only 6 annotations in the Liver group. Inspired by the idea of Group Lasso, however, we propose a group-guided sparsity condition. Specifically, for each SNP *i* and annotation group *g*, we define the group-level annotation as 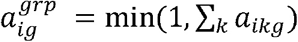. Thus, 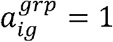 if any annotation *k* in group *g* is positive otherwise zero. In practice, we also included a set of non-tissue-specific baseline annotations, which we impose the same L2-norm regularization as the group-level weights.

At each iteration, we update the group-level enrichments **w**^*grp*^ using RiVIERA-ridge model with the group level annotations. We then set the annotation group *g* to *active* if the corresponding enrichment z-score is greater than a fraction of maximum z-scores among all the group-level annotations *z*_*g*_ > *λ*max(*z*_*g*′_). We then cycle through the active annotation groups and update the corresponding cell-type-specific annotations within each group using Newton-Raphson method, where the Hessian matrix is calculated only within the group. For the inactive annotation groups, we set the corresponding weights of the cell-type-specific annotations to zero. After model converges, we calculate z-score over all of the non-zero coefficients.

In practice, when the enrichment of only a few cell groups (usually fewer than three groups for most traits that we investigated in this paper at fairly lenient sparsity threshold *λ*) are active, this approach is more efficient than the original Group Lasso because at each iteration we do not need to calculate the gradients and Hessian matrix over all of the cell-type-specific annotations but instead only need to calculate the gradients and Hessian matrix of the group-level annotations as well as only the cell-type-specific annotations within each active group.

### 4.4 RiVIERA-fmap fine-mapping model

#### 4.4.1 Likelihood of SNP causal configurations

The proposed fine-mapping algorithm RiVIERA-fmap builds upon some of the existing fine-mapping methods ^17,18,23,26^ by utilizing multivariate normal theory to model the likelihood of the z-scores given a specific *causal configurations I*_*c*_, which is a binary vector with one (zero) indicating the SNP is (not) causal within the locus. We assume that the unobserved effect size **β** follows multivariate normal (MVN) with zero mean and covariance dictated by the phenotypic variance explained by the causal SNPs 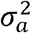 multiplied by the variance of the residual error 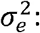 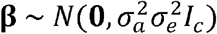 which is assumed in a way that ***β*** can be integrated out.

Specifically, due to conjugate prior of the Gaussian distributions, we can integrate out the effect size **β** to obtain the closed form expression of the z-score likelihood: 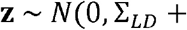 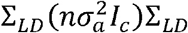, where covariance of the likelihood is a function of the LD matrix ∑_*LD*_, *n* is the effective sample size of the GWAS, and 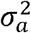 is the variance explained per SNP. In practice the z-score is often estimated by Wald statistics 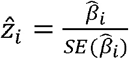 which follows standard normal under the null.

Taking the Bayes factor of the likelihood of a particular causal configuration over the likelihood of the null model of zero causal variant reduces the inference to only the likelihood of the candidate causal SNPs 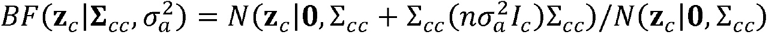. Since we usually do not know 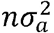, we treat them as a single parameter 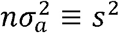. Moreover, instead of inferring a global additive variance, we infer the distribution of the locus-specific variance 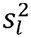 for each locus *l*.

#### 4.4.2 Epigenomic-informed functional priors

We use a logistic function of weighted linear combination of annotations plus the intercepts to infer the prior distribution of each SNP: *π*_*il*_ =*σ*(−_*k*_*w*_*k*_*a*_*ilk*_ +*b*_*i*_), where *σ*(*x*)= 1/(1+ exp(−*x*)) and *w*_*k*_ is the weight for annotation *k* and *a*_*ilk*_ is the annotation *k* for SNP *i* in locus *l*.

The weights **w** = [*w*_1_,*w*_2_,…,*w*_*K*_]′ are interpreted as the “enrichment” parameters. We model the prior distribution of these weights **w** was a K-dimensional multivariate normal distribution over K annotations **w**~*N*(**μ**,*∑*_*w*_) with the *K* × 1 mean vector and the *K* × *K*covariance matrix set to the genome-wide enrichment estimates and the covariance calculated using RiVIERA-glass model (Section 4.2 and 4.3).

The intercepts for each locus *b*_*l*_ captures the belief for each SNP being causal *in the absence of* the annotations (i.e., *π*_*il*_ = *σ*(*b*_*l*_)). We assume *b*_*l*_~*N*(*g*(*π*_0_),1), where the mean as log-logit function *g*(*π*_0_) = log(*π*_0_) − log(1 − *π*_0_) implies that the prior probability of an un-annotated SNP to be causal is 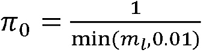, where *m*, is the number of SNPs in the locus and 0.01 is the default (and user defined) maximum prior allowed.

#### 4.4.3 Approximate posterior inference of causal configurations by sampling

The posterior over a configuration **c**_*l*_ for locus *l* is proportional to the product of the likelihood and the prior described above:

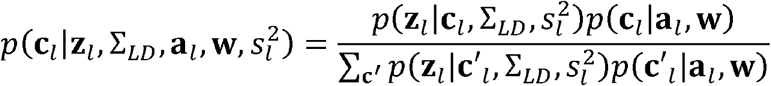

The marginal likelihood in the denominator requires evaluation of exponentially large number of configurations (i.e., 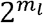 for *m*_*l*_ SNPs in locus *l*), which is intractable. Efficient approximation of the marginal likelihood based on sampling schemes have been proposed in some recent works such as FINEMAP and FastPAINTOR ^18,21^. Inspired by these works, we developed two sampling schemes: (1) Stochastic shotgun sampling by neighborhood search (**Supplementary Note**); (2) Importance Sampling. In our application to the real data, we chose the Important Sampling scheme due to its less reliance on the similarity between the 1000-Genome reference LD matrix and the (unavailable) in-sample LD matrix obtained directly the GWAS individuals, which has profound impact on the summary-statistics-based fine-mapping performance ^61^.

The intuition behind Importance Sampling is that only the configurations containing SNPs with high individual signals in terms of either the marginal SNP z-scores or SNP posteriors are likely to real. Therefore, instead of directly sampling configurations from the actual posterior distribution, which is impractical, we sample from a much easier *proposed distribution* that mimics some of the properties of the desired posterior distribution. In our case, a natural choice of the proposed distribution is the special case in our fine-mapping problem, where each locus contains exactly one causal SNP and a configuration is generated by sampling causal SNPs as independent Bernoulli trials according to their normalized posteriors:

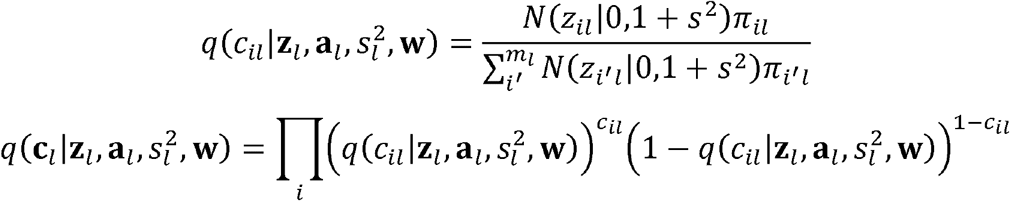

To correct the bias caused by sampling from the proposed distribution, we evaluate the configuration posteriors as follows:

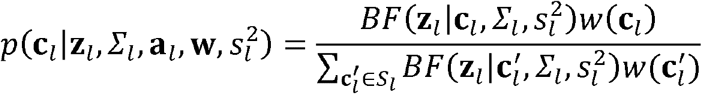

where the importance weight is defined as:

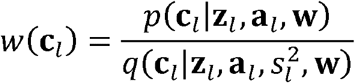

#### 4.4.4 Marginal posterior inclusion probabilities for each SNP

We marginalize these posteriors over all of the configurations *S_L_* to infer posterior inclusion probabilities (PIP) for each SNP: 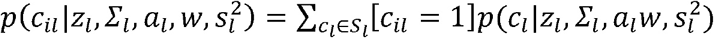. In principle, we would like to infer the expectation of SNP being causal, integrating out all of the model parameters, which is not tractable. As described next, we employ Markov Chain Monte Carlo (MCMC) sampling methods to approximate the integral:

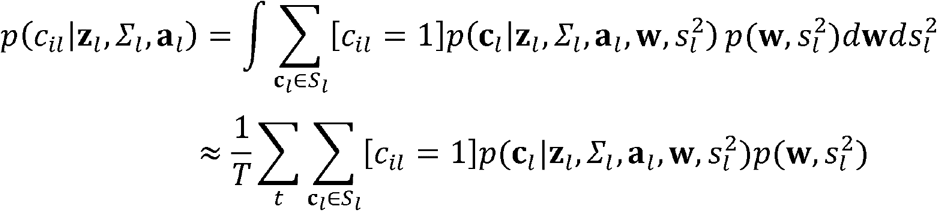

#### 4.4.5 Inferring the distribution of model parameters by Markov Chain Monte Carlo (MCMC)

Given the posterior distribution *q*(***c***_*l*_) of the sampled configurations for each locus *l*, we perform Hamiltonian Monte Carlo (HMC) to sample the model parameters **Θ** = {**w**, **b**, **s**^2^} ^41^. To derive gradients of each model parameters (as required by the HMC method), we define the *expected complete log posterior* as follow:

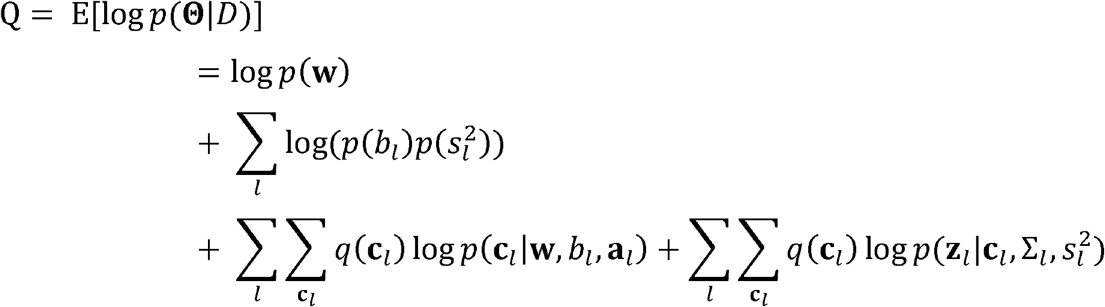

where 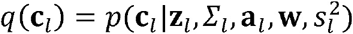 is the posterior distribution over all of the configurations sampled at iteration *t*. Notably, to have a closed-form expression of the objective function, here we assume the z-scores from different loci are conditionally independent given their causal configurations (i.e., the last term). This generally holds true for loci that are far from each other for which we can ensure by merging any loci within 100-kb (or more) genomic distance. The gradients of *Q* with respect to the parameters 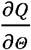 can be easily obtained to enable HMC sampling.

#### 4.4.6 Summary of the RiVIERA-fmap inference algorithm

The unknown parameters are **w**, **b**, **s**^2^, where the enrichment parameters **w** is a *K* × 1 vector for *K* annotations, the intercepts **b** is a *L* × 1 vector for *L* loci, and variance explained per locus **s**^2^ is also a *L* × 1 vector for *L* loci. We summarize the inference algorithm as follow:

1. Initialize the mean and covariance for **w** to be **μ** and *∑*_*w*_ obtained from the genome-wide estimates by RiVIERA-glass, *b*_*l*_ = log(*m*_*l*_) − log(1 − *m*_*l*_) for each locus *l* with *m*_l_ SNPs, and **s**^2^ = **1** for all loci;
2. Given the model parameters **w**, **b**, **s**^2^, infer posterior of causal configurations 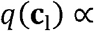 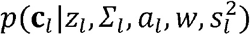 and evaluate and save PIPs 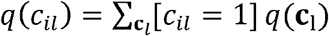 for each SNP *i* in each locus *l*.
3. Given *q*(**c**_l_), jointly sample the model parameters **w**, **b** for all annotations and all loci by HMC;
4. Given *q*(**c**_l_), jointly sample the variance term **s**^2^ for all loci by HMC;
5. If all **w**, **b**, **s**^2^ are accepted at step 3 and 4, do step 2;
6. Repeat step 3 and 5 for *T* MCMC iterations;
7. Estimate PIP for each SNP by averaging over the PIP evaluated at *T*′ < *T* MCMC iterations 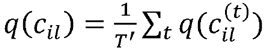.

### 4.5 Preprocessing GWAS summary statistics

Overall, the summary statistics of the 32 GWAS traits were downloaded from public domains at the GWAS consortia website. **Table 1** summarizes the data from each individual GWAS study. For each study, we first removed strand-ambiguous SNP (T/A, C/G) and SNPs with supporting sample sizes lower than a threshold (10,000 individuals). We then imputed summary statistics using ImpG (vl.0.1) (https://github.com/huwenboshi/lmpG) ^27^ to 1000 Genome Phase 1 (version 3) data. Only the imputed SNPs with imputation quality measured as *r*^2^ > 0.6 were retained. Risk loci overlapping with the major histocompatibility complex (MHC) region (chr6:28477797-33448354; hg19) were removed from further analyses. For select risk loci, we calculated the LD for each risk locus by Pearson correlation between the SNPs within each locus using the 1000 Genome European individual-level genotype data.

### 4.6 LD pruning

We ran PriorityPruner (version 0.1.4) (http://prioritypruner.sourceforge.net/) to recursively remove SNPs that are in high LD (*r*^2^ > 0.6) with the most significant SNPs within 100-kb distance. As a result, all of the SNPs from each trait were either labeled as tagged (i.e., pruned) or selected SNPs. We then took only the selected SNPs for the genome-wide enrichment analyses using the proposed RiVIERA-ridge and RiVIERA-glass models described in the main text.

### 4.7 Functional genomic and cell-type-specific Annotations

We annotated each 1000-Genome SNP by the 272 annotations obtained from LD score regression database (https://data.broadinstitute.org/alkesgroup/LDSCORE/) ^16^. There are 52 baseline annotations^62^ and 220 cell-type specific annotations over 10 cell groups. The 220 cell-type-specific annotations were originally generated by the ENCODE and Roadmap Epigenomic Consortium^8^ and represent 100 well defined cell types over four different histone marks namely H3K4me1, H3K4me3, H3K27ac, and H3K9ac.

### 4.8 Transforming LD matrices to invertible matrices

Our fine-mapping method requires the LD matrix to be invertible in order to evaluate the MVN density function and calculate the derivatives of the variance explained **s**^2^. In practice, many LD matrices are not invertible due to finite sample sizes. This posed a problem when fitting our model. To remedy the problem, we transformed all of the LD matrices to invertible matrices using a Kernel-based regularized least squares (KRLS). Similar LD-transformation methods were also proposed elsewhere ^22^. Specifically, we fit a KRLS model one per locus by using the corresponding LD as the predictors and z-scores as the response variable. We implemented the KRLS using the existing R package KRLS ^63^, which finds the best fitting function by minimizing a Tikhonov regularization problem with a squared loss, using Gaussian Kernels as radial basis functions. The original LD matrix is then transformed as *∑*\* = *K* + *λI*. We then further converted the transformed LD matrices to correlation matrices by the R function *cov2cor* to obtain the final transformed LD matrices. The transformed LD matrices have average Pearson correlation > 0.9 with the original LD matrices in terms of the off-diagonal values. Throughout this paper, we used the transformed LD matrices instead of the original LD matrices.

### 4.9 Pathway and GO term enrichment and loss-of-function constraint analysis

Gene set enrichment analysis was performed by hypergeometric test, assessing overrepresentation of genes annotated with GO biological process and canonical pathways in the Molecular Signatures Database (MSigDB)^64^. Gene loss-of-function (LoF)-intolerance scores (pLI scores)^40^. The observed mean pLI score for pleiotropic genes was estimated as the in-sample mean, whereas for non-pleiotropic genes it was estimated by randomly resampling sets of matching size. A null distribution was estimated by randomly resampling with replacement from all scored genes sets of matching size 10,000 times. Significance of deviation from expectation was assessed by computing a z-score and corresponding p-value.

### 4.10 Code availability

RiVIERA software (version 0.9.3) implemented as a standalone R package is freely available from Github repository (https://yueli-compbio.github.io/RiVIERA).

## 5 Acknowledgements

We thank Alkes Price, Hilary Finucane, and Steven Gazal for providing some of the full summary statistics and made publicly available the 272 annotations used in this study. Also, we thank all of the GWAS consortia that made the summary statistics publicly available.

## 6 Author contributions

Y.L developed the method; Y.L. and J.D. analyzed the results. Y.L. wrote the paper with relevant comments and suggestions byJ.D. and M.K.․

